## Supplemental Figure for "Heme’s relevance genuine? Re-visiting the roles of TANGO2 homologs including HRG-9 and HRG-10 in *C. elegans*"

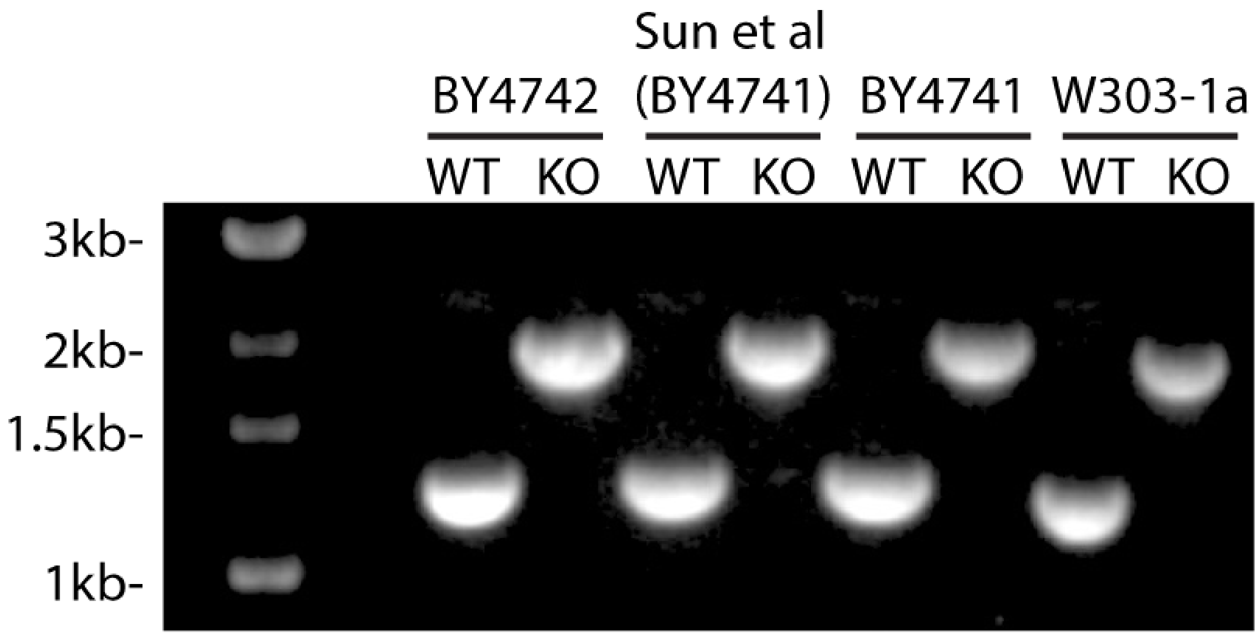


Supplemental Figure 1: PCR confirmation of *YGR127w* knockout cassette integration across yeast strains.
