## Supplementary material for "Heme’s relevance genuine? Re-visiting the roles of TANGO2 homologs including HRG-9 and HRG-10 in *C. elegans*": R Code

Code used data provided by Sun et al. The top 500 differentially expressed genes were plotted and cluster profiling was performed. Data is partially visualized in Figure 1f. Gene ontology was performed using WormCat 2.0 (wormcat.com).

\# Load necessary packages

library(edgeR)

library(gplots)

\# Import the data

gene_data <- read.csv("***.csv", header = TRUE)

\# Extract gene names

gene_names <- gene_data[, 1]

\# Extract count data

condition_counts <- gene_data[, -1]

\# Convert count data to matrix

count_matrix <- as.matrix(condition_counts)

\# Create a DGEList object

dge <- DGEList(counts = count_matrix, genes = gene_names)

\# Normalize and estimate dispersions

dge <- calcNormFactors(dge)

dge <- estimateCommonDisp(dge)

dge <- estimateTagwiseDisp(dge)

\# Design matrix and contrasts (comparing 2 and 400 to 20)

design <- model.matrix(~0 + factor(rep(c("condition20", "condition2", "condition400"), each = 3)))

colnames(design) <- c("condition20", "condition2", "condition400")

\# Define contrasts (comparing 2 and 400 to 20)

contrasts <- makeContrasts(condition2 - condition20, condition400 - condition20, levels = design)

\# Fitting the generalized linear model

dge_fit <- glmFit(dge, design)

\# Likelihood ratio test

dge_contrast <- glmLRT(dge_fit, contrast = contrasts)

\# Get the top differentially expressed genes

top_genes <- topTags(dge_contrast, n = 500)$table

\# Extract expression data for top genes

top_gene_names <- top_genes$genes

top_gene_expression <- count_matrix[which(gene_names %in% top_gene_names), ]

\# Normalize expression data (z-score normalization)

normalized_expression <- scale(top_gene_expression)
#Create distance matrix

dist_matrix <- dist (log2_rnaseq_data, method = “euclidean”)

#Perform hierarchical clustering

Hc_result <- hclust (dist_matrix, method = “complete”)

Num_clusters <- 3

Cluster_cut <- cutree(hc_result, k = num_clusters)
